# Seasonal and directional dispersal behavior in an ongoing dove invasion

**DOI:** 10.1101/517540

**Authors:** David L. Slager

## Abstract

The dispersal behavior that underlies range expansions can be difficult to study. Eurasian Collared-Doves have staged independent northwestward invasions across both Europe and North America, reaching carrying capacity in Europe but continuing to increase exponentially in the Americas, where their dispersal behavior remains enigmatic. I used citizen science observations to investigate Eurasian Collared-Dove dispersal behavior along the North American Pacific coast, a natural barrier to land-bird dispersal. Using coastal and pelagic observations, I assessed the seasonality and directionality of dispersal and its prevalence across years. Dispersal flights peaked in spring, like in Europe, and were north-biased, consistent with northwestward initial colonization. A non-significant trend of increasing dispersal across years may reflect evolution of dispersal via spatial sorting and selection. These results inform management of this invasive dove, raise new questions about evolutionary mechanisms behind the invasion, and exemplify using citizen science to study dispersal behavior, a longstanding challenge in ecology.

## Introduction

Although range expansions of invasive species are readily tracked and modeled using spatial occurrence data (e.g., Gallien et al. 2010, Ingenloff et al. 2017), direct evidence of the dispersal behavior that underlies range expansions is often more elusive. Range expansions obviously require the dispersal of individual organisms, but because dispersal is so difficult to study, we often have a poor understanding of how the movement ecology of dispersal relates to overall colonization and establishment patterns of an invasive species (Ronce et al. 2001, Nathan 2001, Jønsson et al. 2016). Understanding the proximate dispersal behaviors of individuals and groups can help to inform understanding of population-level processes (Macdonald and Johnson 2001). Idiosyncratic dispersal behaviors of species, for example, can aid in predicting distributions and extinctions of populations (e.g., Moore et al. 2008). Moreover, species-specific life history traits, such as asexual reproduction, sperm storage, and simultaneous dispersal of multiple individuals can increase the probability of successful colonization following a dispersal event (Veit and Lewis 1996, Sakai et al. 2001, Duncan et al. 2003, Blackburn et al. 2015).

Replicate invasions of a single species offer a natural opportunity to compare and contrast dispersal behavior while accounting for idiosyncrasies of different invaded environments (Melbourne and Hastings 2009). The Eurasian Collared-Dove (*Streptopelia decaocto*), once restricted to southern Asia, rapidly spread throughout Europe in the 20th century before independently invading the Americas beginning in the late 20th century (Fisher 1953). First introduced into the New World in the Bahamas in the 1970s, Eurasian Collared-Doves colonized Florida by the early 1980s (Smith 1987), reaching the Pacific Coast ~20 years later (Bled et al. 2011). Today, they continue to increase exponentially in western North America at a staggering rate of ~45% per year (Sauer et al. 2017).

The European and American invasions of Eurasian Collared-Doves share some key parallels. First, the initial invasions on both continents progressed primarily northwestward (Coombs et al. 1981, Kasparek 1996, Sullivan et al. 2009). In North America, for example, Eurasian Collared-Doves colonized Alaska prior to the northeastern United States (Sullivan et al. 2009). Second, both colonization histories followed a pattern of “jump” dispersal, with individuals first appearing in far-flung areas and subsequently backfilling intermediate localities (Hudson 1965, Romagosa and Labisky 2000, Fujisaki et al. 2010). Outside of these commonalities, however, we know little about the behavior underlying this dispersal. Furthermore, because of the relatively sparse knowledge of the North American invasion, it is difficult to compare dispersal behavior between the two continents (Romagosa 2012).

Two additional patterns in Eurasian Collared-Dove dispersal behavior have been fairly well characterized in the European invasion but not in the American invasion. Documenting the seasonality of dispersal is important for understanding and modeling biological invasions across the full annual cycle (Hostetler et al. 2015), and although collared-dove dispersal occurred primarily in spring in Europe, little information is available for the Americas (Smith 1987, Kasparek 1996, Romagosa 2012). Second, dispersal intensity of Eurasian Collared-Doves in Europe began to decrease once the species reached carrying capacity there (Marchant et al. 1992, Kasparek 1996). In the Americas, however, where Eurasian Collared-Dove populations are currently increasing exponentially, concurrent trends in dispersal intensity are poorly understood.

Long distance dispersal behavior has historically been challenging and expensive to study. Mark-recapture studies are limited by low re-encounter rates and coarse temporal resolution, global tracking technologies are expensive (Nathan 2001), and genetic markers are seldom informative about behavior at the scale of the individual (Battey et al. 2017). Although tracking technologies and genetic markers continue to improve (Webster et al. 2002, Kays et al. 2015), both approaches continue to be constrained by cost.

Citizen science observations provide an alternative means for documenting animal behavior, especially for highly visible species that occur in populated areas (e.g., Leighton et al. 2018, La Sorte et al. 2018, Freeman and Miller 2018). Moreover, the spatial and temporal breadth of citizen science data may be expedient for overcoming some of the main challenges to studying animal dispersal, an inherently decentralized phenomenon. Eurasian Collared-Doves are a medium-sized, diurnal, social species in which dispersal flights are likely detectable by human observers. Several recent sightings of Eurasian Collared-Dove flocks moving through coastal Washington State hinted at the possibility of dispersal flights, but the anecdotal nature of these observations prevented firm conclusions (DLS pers. obs., RJM, pers. comm.).

To investigate Eurasian Collared-Dove dispersal in North America, I searched for evidence of dispersal-related behavior in a large citizen science database of birding observations. My specific objectives were to 1) document potential seasonality of dispersal-related behavior, 2) characterize the distribution of flight directions during peak dispersal season, and 3) examine potential changes in the frequency of dispersal-related behavior across years.

## Methods

### Citizen science data

To investigate dispersal-related behavior in Eurasian Collared-Doves, I used citizen science observations from the eBird Basic Dataset, a large, worldwide database of birding observations (eBird 2018). I analyzed Eurasian Collared-Dove records from the August 2018 data release, which contained all Eurasian Collared-Dove reports submitted to eBird through August 2018. I conducted all data filtering and analysis in R (R Core Team 2018). Prior to analysis, I discarded observations that failed to meet eBird data quality criteria, and I condensed multi-observer reports to single reports.

### Coastal observations

To focus on a geographic area with numerous citizen science observers and where dispersal behavior was expected to be most visually evident along a major water barrier, I selected observations in the United States and Canada (Alaska, British Columbia, Washington, Oregon, and California) within 8.05 km (5 miles) of the Pacific Ocean (Natural Earth 2018a). Eurasian Collared-Doves tend to be gregarious when not breeding, occurring in flocks of ≥10 individuals, and preliminary anecdotal observations suggested that Eurasian Collared-Doves might conduct long-range movements in flocks (Romagosa 2012; <Redacted>, pers. obs). Thus, I further filtered the dataset to include observations with a count of ≥10 individuals.

To obtain information about dispersal-related behavior in Eurasian Collared-Doves, I selected records that included a non-empty text string (“species comments”) in which observers are able to enter field notes about a species observation. To code behavior, I manually parsed these descriptions for each record while blind to other fields. I considered descriptions of Eurasian Collared-Dove flocks in directional flight to represent instances of potential dispersal-related behavior. I conservatively coded records as describing a “flying flock” when the description explicitly mentioned ≥2 birds flying together, excluding records that explicitly mentioned birds flushing (e.g., from a predator) or traveling to or from a roost site. I coded descriptions of flight directions as the eight cardinal and intercardinal directions, circular/out-and-back, or NA. I used coastal observations to investigate seasonality, flight direction, and trends in dispersal-related behavior across years.

### Pelagic observations

Eurasian Collared-Doves are such accomplished dispersers that during the European invasion they occasionally landed on ships in the Atlantic Ocean (e.g., Casement 1983). With this in mind, and to measure dispersal phenology independent of land-based observations, I also queried the eBird Basic Dataset for pelagic sightings of Eurasian Collared-Doves off the coast of western North America. I selected Pacific Ocean locations >8.05 km (>5 miles) from land in the western United States and Canada (Natural Earth 2018b). I manually filtered the dataset, removing records from small islands not recognized as land in the Natural Earth shapefile, removing a few erroneously plotted terrestrial sightings, and removing records from long boat transect surveys that appeared to include both terrestrial and pelagic observations. As an additional quality control step and to avoid duplicate records, I sorted observations by date and time, scanning observer comments to conservatively eliminate reports potentially made by different observers on the same vessel and reports potentially involving collared-dove(s) riding along on a vessel. Unlike with the land-based coastal dataset, I considered all pelagic occurrences of Eurasian Collared-Doves to represent potential instances of dispersal, and did not filter pelagic observations for flock size or code flight behaviors. I used pelagic observations to investigate seasonality only.

### Observer effort

I obtained effort information from the prepackaged Sampling Event Data file from the October 2018 release of the eBird Basic Dataset. To ensure consistency with the Eurasian Collared-Dove dataset, I excluded checklists from after August 2018 and condensed multi-observer checklists to single checklists.

As an index of coastal observer effort, I used a custom R script to quantify the number of terrestrial birding outings (“eBird checklists”) submitted to Project eBird within 8.05 km (5 miles) of the Pacific Ocean in the western United States and Canada. For an index of pelagic observer effort, I used the number of boat-based eBird checklists >8.05 km (>5 miles) from land off the western United States and Canada.

### Seasonality

To assess seasonality in coastal reports of flying flocks, I conducted a *χ*^2^ test of the raw number of reports of flying flocks each month, simulating p-values with 2,000 Monte-Carlo replicates (Hope 1968). To control for potential monthly variation in observer effort, I also conducted a *χ*^2^ test with expected monthly probabilities set to monthly proportions of coastal observer effort.

Similarly, to examine the seasonality of pelagic observations, I conducted one *χ*^2^ test on the raw monthly number of reports, and a second *χ*^2^ test using monthly proportions of pelagic observer effort as the expected monthly probabilities.

### Flight direction

To characterize the flight direction of coastal collared-dove flocks exhibiting dispersal-related behavior during peak dispersal months of March-May, I calculated the mean flight direction 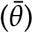, mean resultant length, circular variance (1 - mean resultant length), and directional range using R package circular (Pewsey et al. 2013, Agostinelli and Lund 2017) after removing records with a NA or circular/out-and-back flight direction. I tested the null hypothesis of uniform flight direction against an alternative hypothesis of a due north mean flight direction using a Rayleigh Test (Jammalamadaka and Sengupta 2001). For flight directions outside of the months of March-May, I used a Rayleigh Test to determine if the distribution of flight directions differed significantly from a radially uniform distribution.

### Frequency of dispersal-related behavior across years

To assess trends in the frequency of collared-dove spring dispersal across years, I conducted a linear regression with year as the explanatory variable, including all years with spring coastal reports of Eurasian Collared-Dove flying flocks (2010-2018). To control for increases in collared-dove populations and increases in citizen science participation across years, I set the response variable to the number of March-May flying flock observations divided by the total number of March-May Eurasian Collared-Dove reports.

### Qualitative observations

To gauge how citizen science observers interpreted their own Eurasian Collared-Dove sightings, I scanned the text comments for records involving flying flocks. I noted instances when observers speculated about dispersal or migration and when observers commented that the behaviors seemed novel or unusual.

### Data accessibility

Coded observations and R scripts for filtering, analyzing, and visualizing the data are available on GitHub at https://github.com/slager/collared_dove_dispersal and will be posted on Dryad or a similar archive prior to final publication.

## Results

### Data filtering

The August 2018 release of the eBird Basic Dataset contained 1,862,997 Eurasian Collared-Dove reports. Collapsing multiple observer reports to single reports and limiting the dataset to Pacific coastal localities in the western United States and Canada yielded 1,997,921 eBird checklists, including 152,420 Eurasian Collared-Dove observations from 9,043 observers at 21,538 coastal localities. Additional filtering to reports of ≥10 individuals with text comments resulted in 841 records from 336 observers at 440 coastal localities. Among these 841 records, flying flocks were described in 95 reports from 56 observers at 65 localities (Figure 1, Figure 2). Across all months, 32 of 95 reports of flying flocks (34%) specified a flight direction, including two noted as circular or out-and-back.

**Figure 1:**
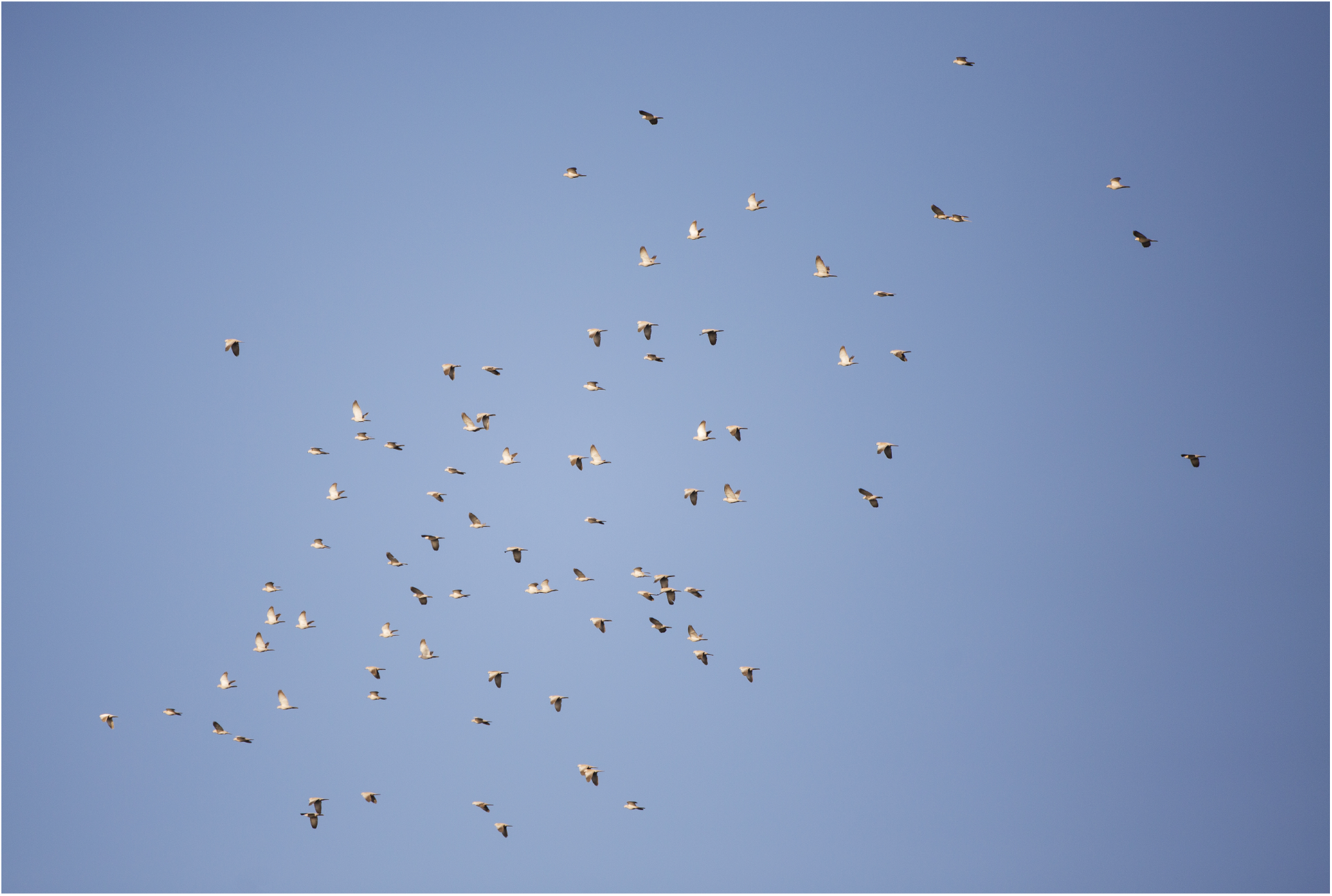
Eurasian Collared-Doves in flight over the Puget Sound at Point No Point, Washington on 26 April 2018. Photo © Janine Schutt, used with permission.

**Figure 2:**
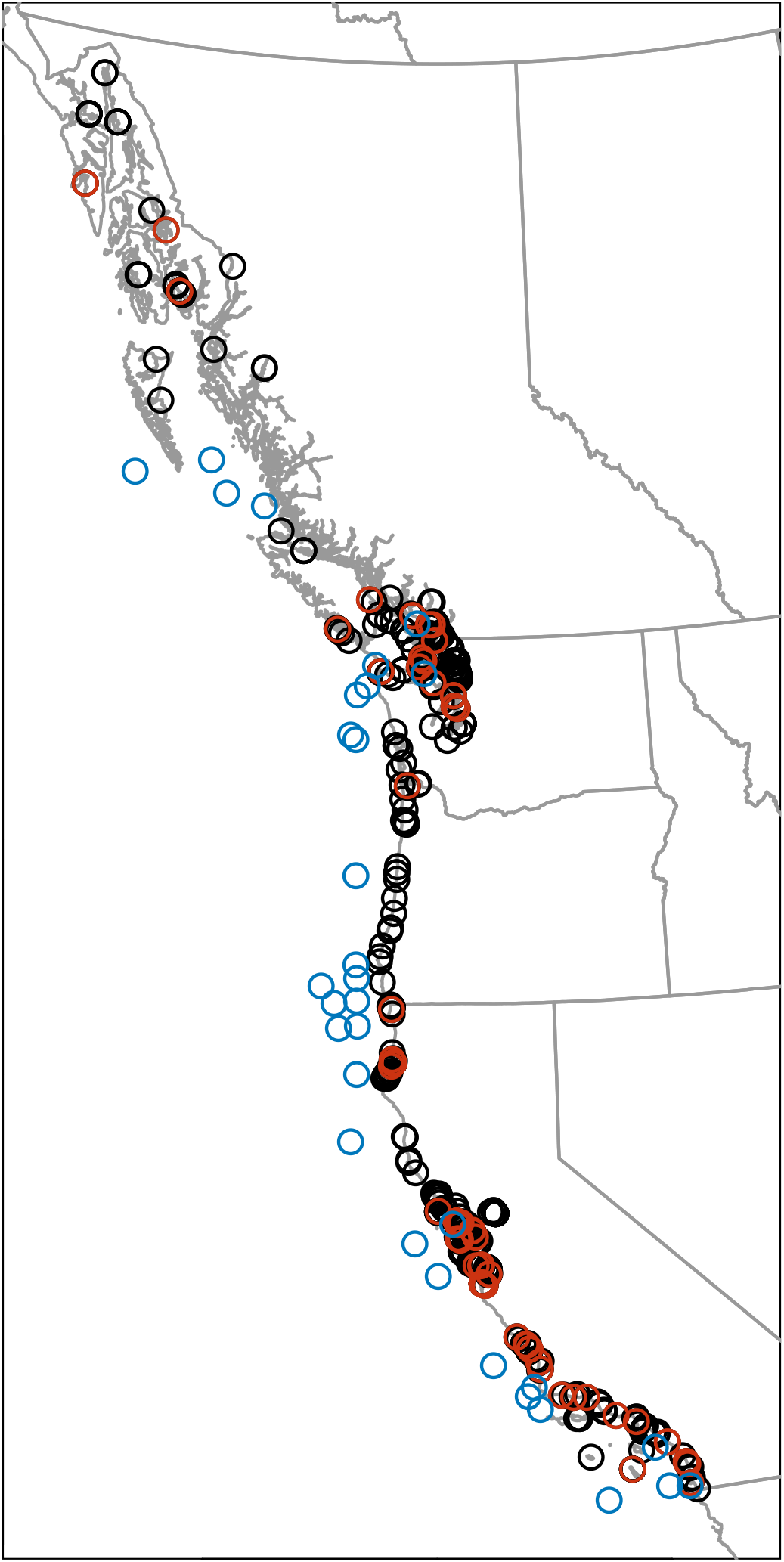
Locations of Pacific coastal observations of ≥10 Eurasian Collared-Doves with field notes (black; n=841), the subset of these observations that described flying flocks (red; n=95), and locations of pelagic Eurasian Collared-Dove reports (blue; n=32).

The final pelagic dataset of locations in the Pacific Ocean offshore from the western United States and Canada contained 28,012 eBird checklists, with 32 records of Eurasian Collared-Doves from 23 observers at 32 localities (Figure 2).

### Seasonality

A majority of coastal reports of flying flocks (56 of 95, or 59%) occurred during the months of March, April, and May (Figure 3A, *χ*^2^ = 82.22, p < 0.0005). This seasonality persisted when controlling for observer effort (55%, *χ*^2^ = 62.85, p < 0.0005). Flying flocks were reported more often during each of the months of March, April, and May than in any of the other 9 months of the year, with and without controlling for observer effort.

**Figure 3:**
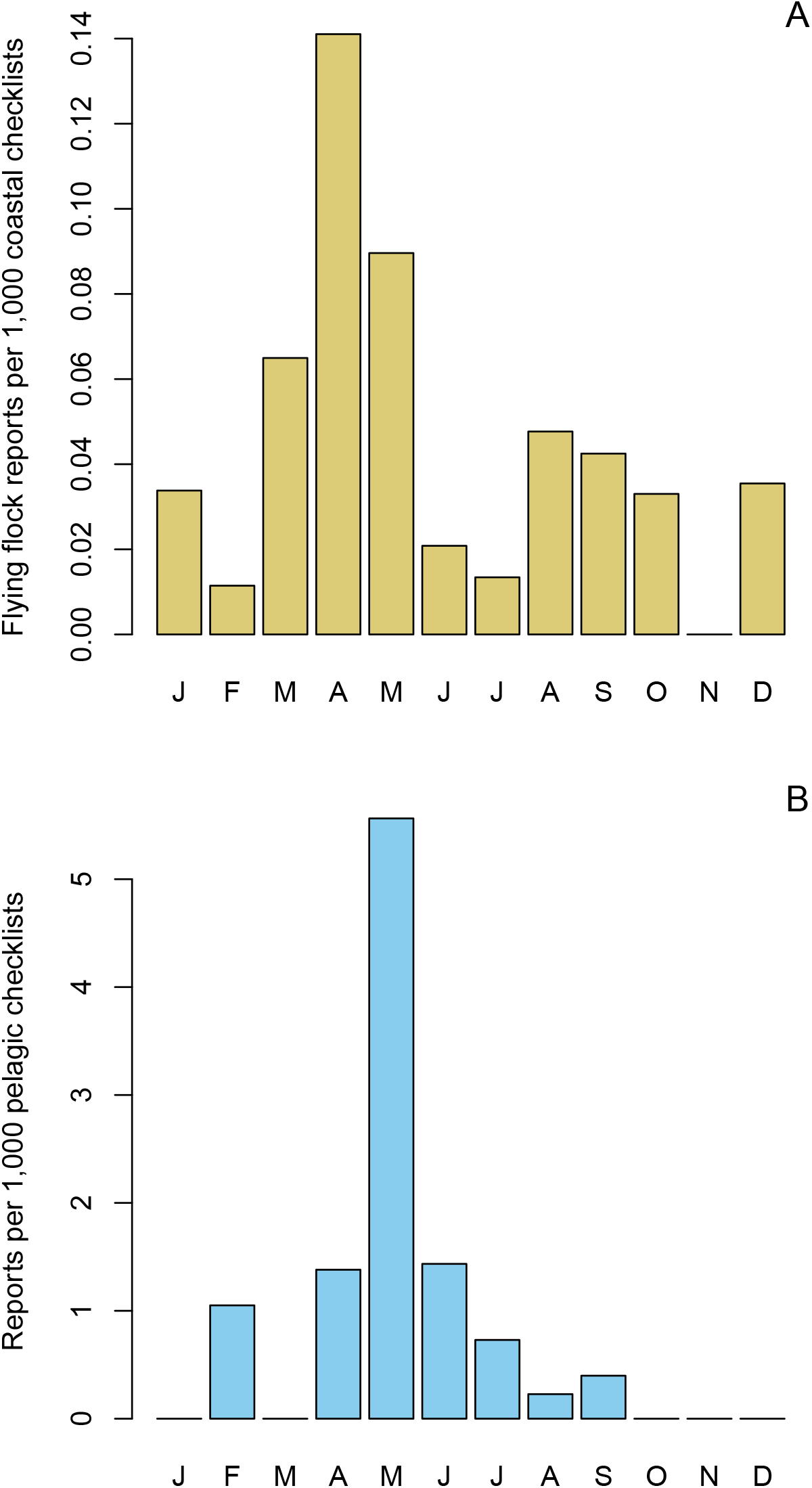
Seasonality of A) Pacific coastal Eurasian Collared-Dove flying flocks and B) pelagic Eurasian Collared-Dove sightings off the Pacific coast of the United States and Canada.

Pelagic records were also concentrated during the spring season, with 23 (72%) occurring during the months of March-May, including 20 (63%) in May alone (Figure 3B, *χ*^2^ = 128.5, p < 0.0005). This pattern was also present after controlling for monthly variation in observer effort (64% March-May; 52% May only; *χ*^2^ = 75.89, p < 0.0005).

### Flight direction

During the March to May peak season for reports of flying flocks, 26 of 56 observations of flying flocks (46%) specified a flight direction, including one described as circular or out-and-back. The remaining 25 that specified a compass direction were made by 15 observers at 17 localities. The distribution of these March-May flight directions was non-uniform (Figure 4, Rayleigh Test, test statistic = 0.897, p < 1×10^−8^). Most flocks (68%) were flying due north, and flight directions were north-biased (Rayleigh Test, *μ* = 0°, test statistic = 0.878, p < 1×10^−9^). The mean flight direction was 348°, the mean resultant length was 0.897, and the circular variance was 0.103. The range of flight directions spanned 135° from due west via north to northeast.

**Figure 4:**
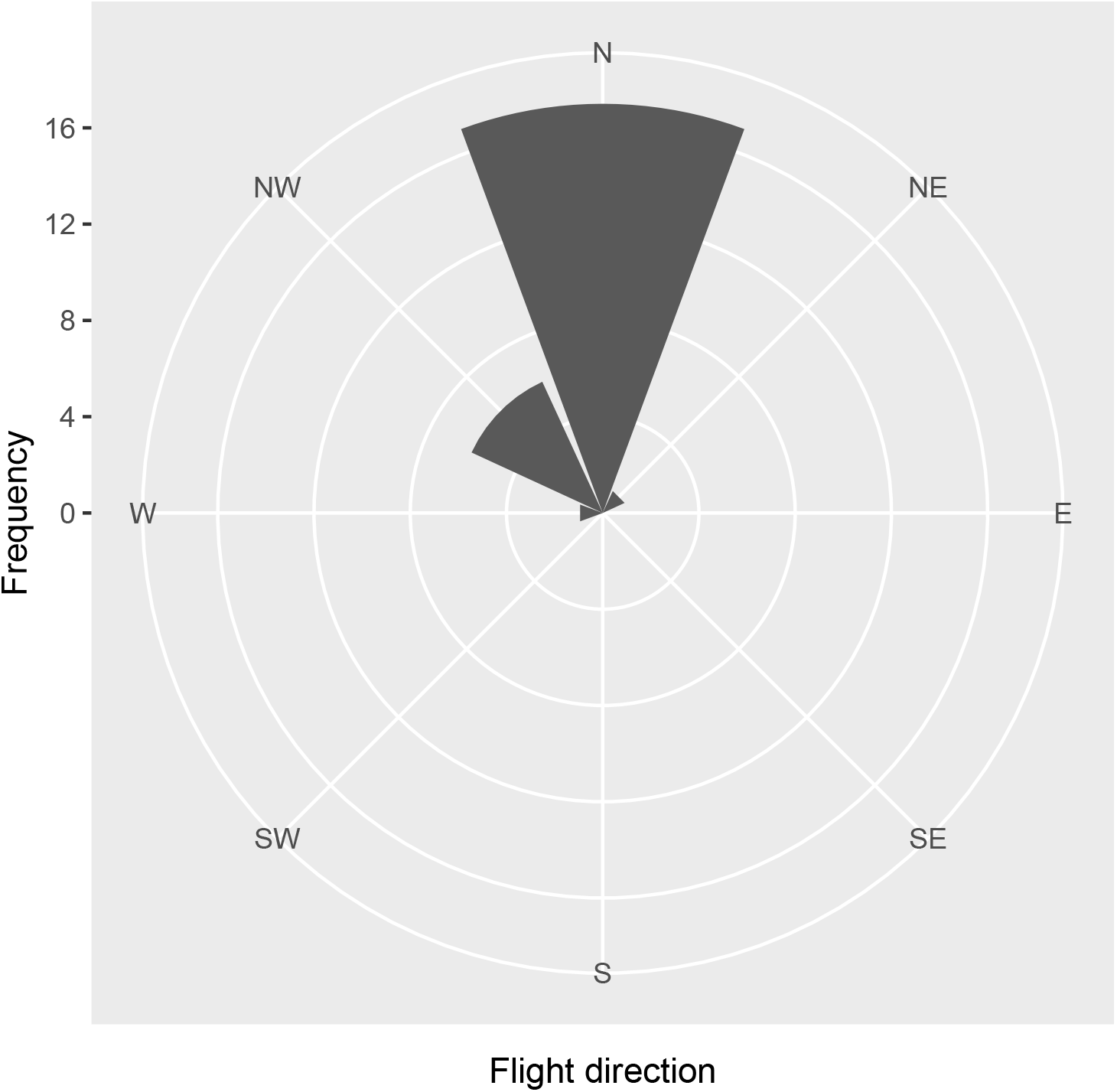
Flight direction of March-May flocks in reports of ≥10 Eurasian Collared-Doves along the Pacific coast of the United States and Canada.

Outside of the months of March-May, 5 of 39 observations of flying flocks (13%) specified a flight direction, including 3 northbound, 1 westbound, and 1 eastbound. This distribution of flight directions did not differ from uniform (Rayleigh Test, test statistic = 0.6, p = 0.17).

### Frequency of dispersal-related behavior across years

After controlling for annual increases in the number of Eurasian Collared-Dove sightings, there was a non-significant trend of increasing spring frequency of flying flock observations across years (F_1,7_ = 1.82, R^2^ = 0.09, p = 0.22). The trend was consistent with an 85% increase in the frequency of spring flying flocks from 2010-2018, or a mean geometric increase of 8% per year.

### Qualitative observations

Of the 95 observations that described flying flocks in text comments, eight (8%) described these behaviors as novel or unusual. Eleven of the 95 reports speculated that the birds observed were migrating or dispersing, and of those 11 observations, ten (91%) took place in March-May.

## Discussion

Aggregate citizen science observations of Eurasian Collared-Dove behavior indicate that Pacific coastal dispersal flights peak during spring and are north-biased. Furthermore, these flights have become detectable in eBird data only relatively recently (since 2010), and show a non-significant increase in frequency across years. Although no data is available on the cohesion or final destinations of spring flying flocks along the Pacific Coast, flocking dispersal behavior presents a ready mechanism for “jump” dispersal in Eurasian Collared-Doves. By dispersing in flocks, collared-doves may circumvent Allee effects like the demographic time lags, density thresholds, and stochastic vulnerabilities that disproportionately affect solitary dispersers (Courchamp et al. 1999, Stephens and Sutherland 1999, Simberloff 2009, Schmidt et al. 2015, Battey 2018).

### Spring dispersal, natal dispersal, and the annual cycle

Dispersal-related behavior in Eurasian Collared-Doves peaked in March-May both along the Pacific coastline and in pelagic waters. The May pelagic peak occurred later than the coastal April peak, but this difference could be an artifact of low pelagic sample size and/or clumped timing of pelagic birding trips and northbound repositioning cruises.

Peak dispersal of Eurasian Collared-Doves along the Pacific Coast in spring is consistent with the seasonality of dispersal during the European invasion, where sightings also peaked during spring on the North Sea island of Helgoland (Kasparek 1996) and on Atlantic Ocean ships (Smith 1987). Mark-recapture data from collared-doves in Europe showed a rapid increase in average ringing recovery distance during the spring of a collared-dove’s second calendar year (Kasparek 1996), consistent with spring dispersal and also suggesting a prominent role for natal dispersal. Elevated natal dispersal relative to adult dispersal occurs in birds generally (Winger et al. 2018) and in collared-doves specifically (Coombs et al. 1981, Baptista et al. 1997), where some natal dispersal events can exceed 1000 km (Baptista et al. 1997).

Spring natal dispersal makes sense in the context of the full annual cycle of Eurasian Collared-Doves. In North America, hatch-year birds complete their first prebasic molt in April-November near their natal area or on their non-breeding grounds, and molt can be slowed, suspended, or arrested in November-March (Romagosa 2012). Eurasian Collared-Doves can breed in their second calendar year, and peak breeding (across all ages) occurs in June-August, even though some individuals breed earlier (Coombs et al. 1981). Thus, the March-May natal dispersal period occurs after molting and overwintering but prior to peak breeding and the onset of the definitive prebasic molt in April-October (Romagosa 2012). Additional investigation of the age structure of dispersing Eurasian Collared-Doves relative to the age structure of the overall population is needed to establish the ontogeny and prevalence of natal dispersal in North America (Massot et al. 2002).

### Dispersal vs. seasonal migration

North-biased spring movements of Eurasian Collared-Doves along the Pacific coast of North America could be interpreted as either dispersal or seasonal migration, but several lines of evidence strongly favor dispersal. First, spring dispersal was well documented in the European invasion (Kasparek 1996). Second, although some high-elevation south Asian populations of Eurasian Collared-Doves are partially migratory (Fisher 1953), the species is considered non-migratory in North America (Romagosa 2012). Finally, if spring flights along the Pacific Coast represented seasonal migration, then we might expect to observe a corresponding fall migration. However, no signal of fall migration was evident from citizen science observations. Although my investigative approach could have missed fall migrants if Eurasian Collared-Doves in autumn migrate singly and/or take a more inland route, I am not aware of any evidence for such movements.

One way to directly differentiate between seasonal migration and dispersal would be to investigate Eurasian Collared-Dove movement ecology using modern tracking devices (e.g., Kays et al. 2015). However, future citizen science data might also provide a means to test this question. If spring Eurasian Collared-Dove movements along the Pacific Coast represent spring migrants, then these spring movements should increase concomitantly with growing northern populations of collared-doves and persist into the future. On the other hand, if spring movements represent dispersal, then spring flight directions should become more uniform once collared-dove populations reach carrying capacity throughout North America.

### Flight direction

Spring flocks of Eurasian Collared-Doves had strongly north-biased flight directions. This flight direction is consistent with the northwestward vector of initial colonization in both the American and European invasions. This is because northwestward-dispersing birds from inland populations would be expected to accumulate upon reaching the Pacific Ocean and deflect northward along the coast. Although northbound flocks along the Pacific coast are also consistent with birds of coastal origin moving due north, coastal birds represent a much smaller potential source population than inland birds, and thus likely constitute only a minority of individuals in northbound coastal flocks. One potential concern with citizen science flight direction data could be that observer presuppositions about spring migration might lead to north-biased reports of spring flight direction when actual spring flight directions are more uniform. However, if this were the case, we would also expect to detect an apparent increase in reports of southbound flocks during the fall months, which was not evident in the data.

### Frequency of dispersal-related behavior across years

After controlling for increases in the overall number of collared-dove reports due to ongoing rapid increases in both collared-dove populations and citizen science participation, reports of spring flying flocks of Eurasian Collared-Doves showed a non-significant increase in frequency across years. Although the trend was not statistically significant, the overall pattern observed is consistent with processes like spatial sorting and selection driving an increase in dispersal behavior along an expanding range front (Lindström et al. 2013, Brown et al. 2014, Ochocki and Miller 2017). The evolution of dispersal behavior in Eurasian Collared-Doves merits further mechanistic study, given rapid northwestward dispersal across two continents from a relatively stable initial distribution in southern Asia (Fisher 1953). Continued monitoring of dispersal behavior is needed to determine whether dispersal will wane once the species reaches carrying capacity in North America, as happened in Europe (Marchant et al. 1992, Kasparek 1996). A decrease in dispersal behavior after population expansion slows is consistent with relaxed selection for dispersal, selection against dispersal, and/or the dissipation of spatial sorting (Marchant et al. 1992). Eurasian Collared-Doves in North America represent an excellent opportunity to study an expanding range edge, and collecting and archiving genetic samples before, during, and after the period of exponential expansion will enable researchers to investigate selection on loci related to dispersal behavior, allele surfing (e.g., Klopfstein et al. 2006), and local adaptation (e.g., Bay et al. 2018).

### Strengths and limitations of inferences from citizen science data

Inferring behavioral patterns from citizen science field notes comes with some caveats. Although I used reports of flying flocks as a metric for detecting dispersal in Eurasian Collared-Doves, I do not claim that all these reports of flying flocks represent dispersal events, and local flock movements certainly contributed to noise in the data. Remarkably, despite this noise, strong patterns of seasonality and directionality were evident in the dataset. Moreover, these patterns were supported by observations from many citizen science observers at many different locations. These quantitative conclusions were also corroborated by qualitative comments by multiple eBird observers noting recent changes in Eurasian Collared-Dove behavior at their regular birding locations. Overall, only a small proportion of Eurasian Collared-Dove observations I analyzed contained text comments, suggesting that the majority of spring dispersal flights of Eurasian Collared-Doves simply go unnoticed or unreported.

## Conclusion

Citizen science provides spatially and temporally decentralized observations of animal behavior at scales that were formerly impractical to study. By examining citizen science field notes in Eurasian Collared-Dove observations along the Pacific Coast water barrier of North America, I was able to measure the seasonality, directionality, and frequency of dispersal across years in this species while it rapidly invades western North America. These spring dispersal flights documented by citizen scientists likely represent the proximate mechanism by which Eurasian Collared-Doves are expanding their range northwestward across North America. Firmly placing Eurasian Collared-Dove dispersal within the spring portion of the annual cycle has important implications for full life cycle modeling and management of this recent arrival to North America (Reed 1999, Macdonald and Johnson 2001, Bowler and Benton 2005). Overall, the phenology, direction, and potentially increasing occurrence of dispersal flights over time closely mirror the European invasion several decades ago. This new understanding of collared-dove dispersal provides a starting point for investigating potential mechanisms for changes in dispersal behavior over time as Eurasian Collared-Doves in North America transition from the exponential growth to carrying capacity.

## Acknowledgments

I thank the many citizen scientists on the Pacific coast of North America who described Eurasian Collared-Dove behavior in their eBird reports. K. Epperly, C. French, J. Klicka, E. Linck, J. Marzluff, S. McNew, R. Merrill, and S. Rohwer provided helpful comments and discussion that improved the manuscript.

